# MiroSCOPE: An AI-driven digital pathology platform for annotating functional tissue units

**DOI:** 10.1101/2025.07.11.664228

**Authors:** Madeleine R. Fenner, Selim Sevim, Guanming Wu, Deidre Beavers, Pengfei Guo, Yucheng Tang, Christopher Z. Eddy, Kaoutar Ait-Ahmad, Travis Rice-Stitt, George Thomas, M.J. Kuykendall, Vasilis Stavrinides, Mark Emberton, Daguang Xu, Xubo Song, S. Ece Eksi, Emek Demir

## Abstract

Cancer tissue analysis in digital pathology is typically conducted across different spatial scales, ranging from high-resolution cell-level modeling to lower-resolution tile-based assessments. However, these perspectives often overlook the structural organization of functional tissue units (FTUs), the small, repeating structures which are crucial to tissue function and key factors during pathological assessment. The incorporation of FTU information is hindered by the need for detailed manual annotations, which are costly and time-consuming to obtain. While artificial intelligence (AI)-based solutions hold great promise to accelerate this process, there is currently no comprehensive workflow for building the large, annotated cohorts required. To remove these roadblocks and advance the development of more interpretable approaches, we developed MiroSCOPE, an end-to-end AI-assisted platform for annotating FTUs at scale, built on QuPath. MiroSCOPE integrates a fine-tunable multiclass segmentation model and curation-specific usability features to enable a human-in-the-loop system that accelerates AI annotation by a pathologist. The system is used to efficiently annotate over 71,900 FTUs on 184 prostate cancer hematoxylin and eosin (H&E)-stained tissue samples and demonstrates ready translation to breast cancer. Furthermore, we publicly release a dataset named Miro-120, consisting of 120 prostate cancer H&E with 30,568 annotations, which can be used by the community as a high-quality resource for FTU-level machine learning aims. In summary, MiroSCOPE provides an adaptable AI-driven platform for annotating functional tissue units, facilitating the use of structural information in digital pathology analyses.

## Introduction

Within an organ, cells organize into functional tissue units (FTUs): small, repeating structures performing essential functions such as nephrons, alveoli, or glands^1^. Changes to these structures, visible in hematoxylin and eosin (H&E)-stained tissue, offer critical information regarding the diagnosis and treatment of multiple diseases, including cancer. Driven by the increased digitization of tissue samples and advances in artificial intelligence (AI), there has been a rapid expansion of digital pathology approaches to harness the detailed biological information in tissue images for improved disease characterization and prediction. These approaches vary in their level of granularity, with some viewing the tissue as a collection of image tiles^2–5^, while others zoom in to the single-cell level to examine cellular interactions and neighborhoods^6–9^, often utilizing data from immunofluorescence or omics techniques. At each extreme, context around tissue structure is largely lost; tile-level methods arbitrarily cut through FTUs, disrupting their morphological arrangement, while cell-level methods fail to capture broader spatial organization. This disconnect in granularity makes it challenging to relate model representations to clinical knowledge, limiting interpretability and posing challenges for incorporating feedback from human experts. This, in turn, creates serious barriers to widespread clinical trust and adoption.

The limiting factor for incorporating FTU level information is the need for detailed annotation by experts, a costly process that demands significant time and manual effort. Leveraging AI to automate the annotation process is critical for reaching the scale of annotated images needed. There are several existing AI models for segmenting and classifying tissue structures in various organs: Many annotate at the regional level, where groups of FTUs of a similar pattern are segmented together within areas that include the intervening stroma^10–12^. While regional annotations provide an overview of tissue arrangement, they lack the precision required to accurately represent intratumoral heterogeneity and model individual FTU features, distributions, and interactions within the tumor microenvironment. Models that perform true FTU-level annotation are rare, and few attempt segmentation and classification for all FTU structures, covering non-tumoral, tumoral, and stromal components^13–16^. Additionally, applying existing models for custom research aims and new datasets presents many challenges. Even when models are trained with diverse data to enhance generalizability, distribution shifts due to variations in tissue collection or processing methods can lead to drops in performance^17^. Most models are offered as static tools without the infrastructure for fine-tuning, limiting adaptability when zero-shot inference underperforms. Furthermore, dataset curation processes are often *ad hoc*, lacking integration with widely adopted annotation viewing and editing platforms and standardization of file types, ontologies, and annotation procedures.

To address these challenges, we developed MiroSCOPE, a platform for annotating FTUs, which employs a human-in-the-loop (HITL) approach to enable large-scale curation by an expert (Figure 1). We combined a fine-tunable AI model for segmentation and classification with a QuPath-based user interface that is optimized to rapidly annotate FTUs. We demonstrate the improvements in efficiency and accuracy offered by the platform using 184 prostatic acinar adenocarcinoma H&E images sourced from three distinct cohorts, annotating over 71,900 FTU structures with uniquely detailed labeling of non-tumoral, tumoral (with subpatterns), and stromal components. We publicly release a subset of these annotated images as a high-quality dataset named Miro-120 (120 prostatic acinar adenocarcinoma H&E; 30,568 annotations). The dataset provides a novel level of structural detail for prostate cancer tissue, and may act as a resource for new structurally informed machine learning efforts.

**Figure 1.**
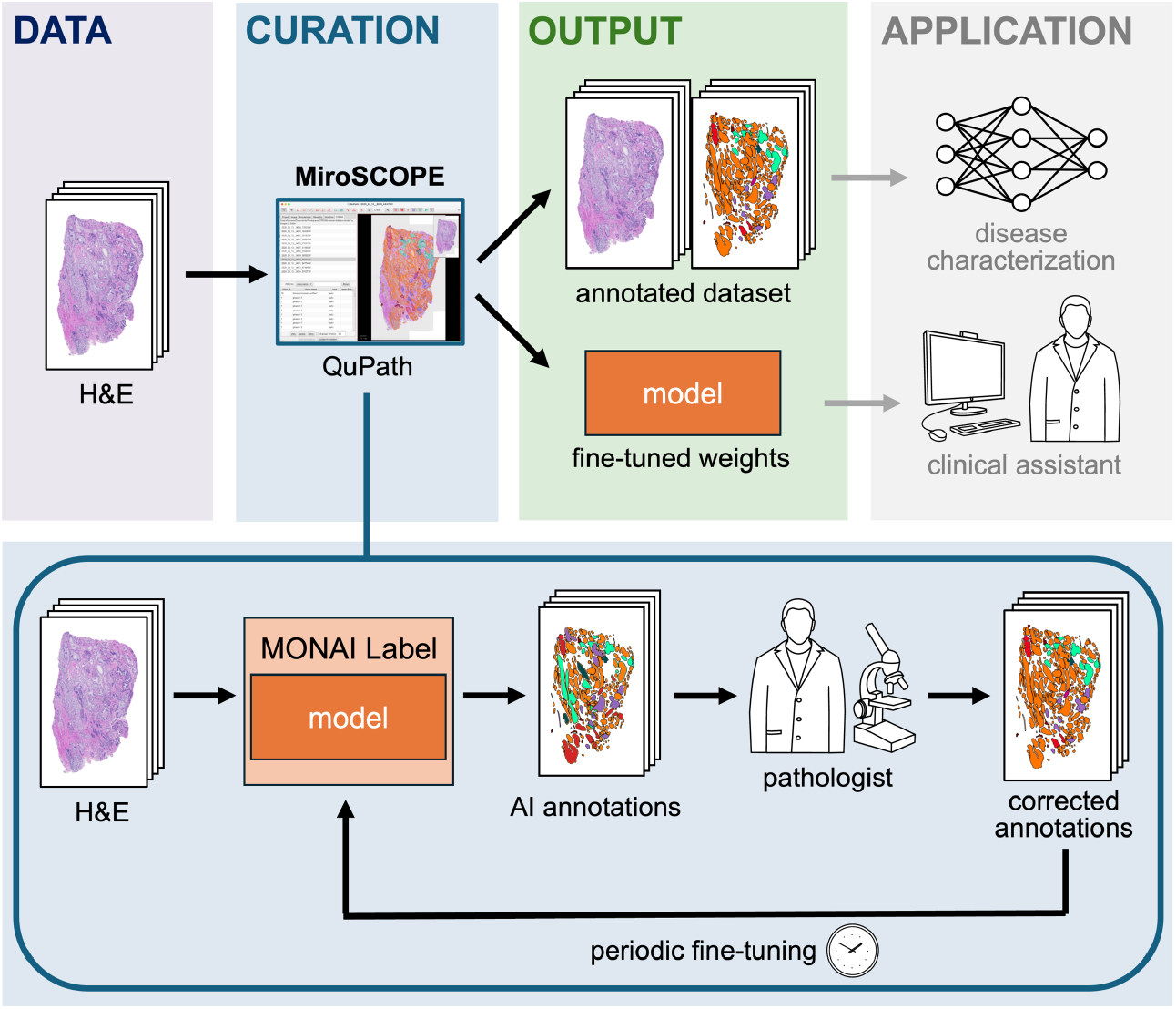
Workflow for employing MiroSCOPE. MiroSCOPE is a QuPath extension that enables the large-scale annotation of FTUs on H&E images. Using a MONAI Label framework, the extension integrates a SAM-based annotation model for automatic segmentation and classification and also offers user interface features specific to the curation task. By enabling a human-in-the-loop approach with expert corrections by a pathologist, a high-quality annotated dataset and fine-tuned model are generated. These outputs can be valuable assets for downstream applications, including disease characterization modeling and clinical tool development.

Prostate cancer is an ideal use case for the system; despite high rates of early diagnosis, a significant subset progresses to advanced and incurable disease, highlighting the need for improved disease staging and prognostic tools^18^. Additionally, prostate cancer is largely characterized by changes in the architectural arrangement of glandular FTUs, but the Gleason scoring system used to categorize this change faces high interobserver variability^19–21^. Employing AI-based approaches to characterize tissue structures holds great promise to reduce variability and increase the prognostic accuracy for prostate cancer and for many other cancer types. We showed the adaptability of the platform to other cancers using a small breast adenocarcinoma dataset in a *few-shot* setting.

## Results

### A controlled vocabulary for annotating prostate cancer FTUs

There is no formal ontology for describing the different types and presentations of FTUs in the disease state. We designed a simple, 17-term controlled vocabulary to annotate FTUs of the prostate for tissue samples with prostatic acinar adenocarcinoma, encompassing diverse histopathologic patterns. The FTUs can be divided into two major categories: glandular origin and stromal. The FTUs of glandular origin cover non-tumoral gland structures, defined as either normal or atrophic, as well as gland-derived tumoral structures. The tumoral FTU classes of the vocabulary follow the Gleason scoring system^19^, the standard metric for evaluating prostate tumors based on the architectural arrangement of glands, ranging from well-differentiated (Gleason Pattern 3) to poorly differentiated (Gleason Pattern 5). Gleason patterns can be subtyped based on morphological characteristics, and these divisions are captured in the labeling scheme. Prostatic intraepithelial neoplasia (PIN) is also included as a precursor. Stromal components, which may provide valuable insights into tumoral behavior, include nerves, vascular structures, and areas of inflammation. Air bubbles, tissue folding, unfocused scanning areas, color anomalies related to tissue processing, and, most prevalently, sheared pieces of glandular tissue, which cannot be evaluated, were annotated as artifacts. The categories of FTUs are summarized in Figure 2.

**Figure 2.**
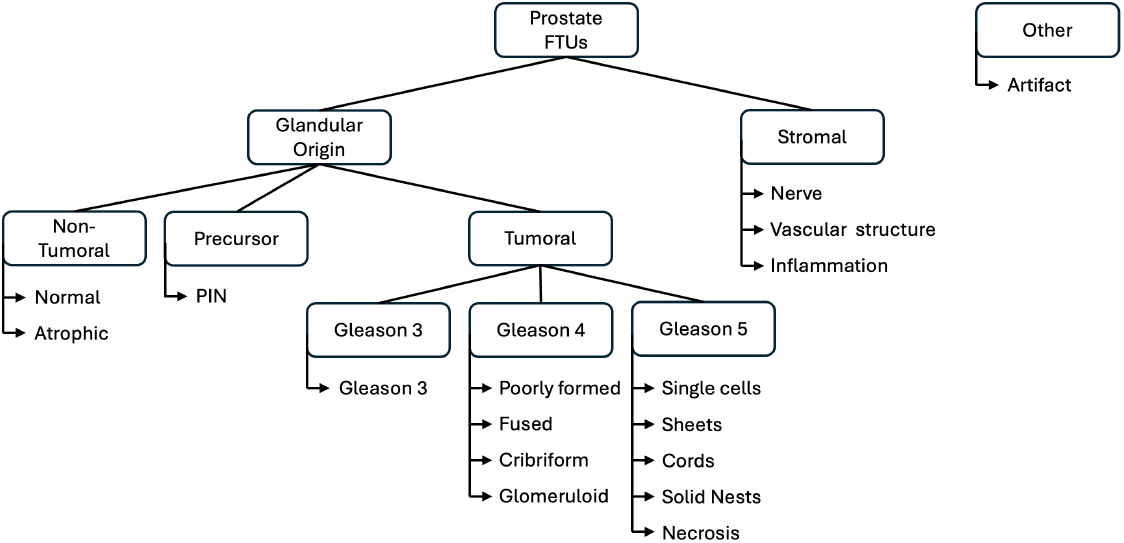
Hierarchical organization of prostate FTU categories defined by the controlled vocabulary. The controlled vocabulary contains 17 terms covering FTUs observed in prostate tissue, with tumoral labeling focusing on prostatic acinar adenocarcinoma. PIN: prostatic intraepithelial neoplasia.

### MiroSCOPE: A platform for human-in-the-loop annotation at scale

To enable the large-scale curation of FTU annotations, we built a HITL system in which a pathologist iteratively uses and improves an AI annotation model. The software stack uses: (i) QuPath^22^, a widely used open-source software for bioimage visualization and analysis, as a base platform, (ii) an AI model fine-tuned to segment and classify FTUs (iii) MONAI Label^23^ to bridge the AI model to QuPath, and (iv) the MiroSCOPE extension, developed by our team to integrate AI inference capabilities and add additional user interface (UI) features specific to the curation task (Figure 3).

**Figure 3.**
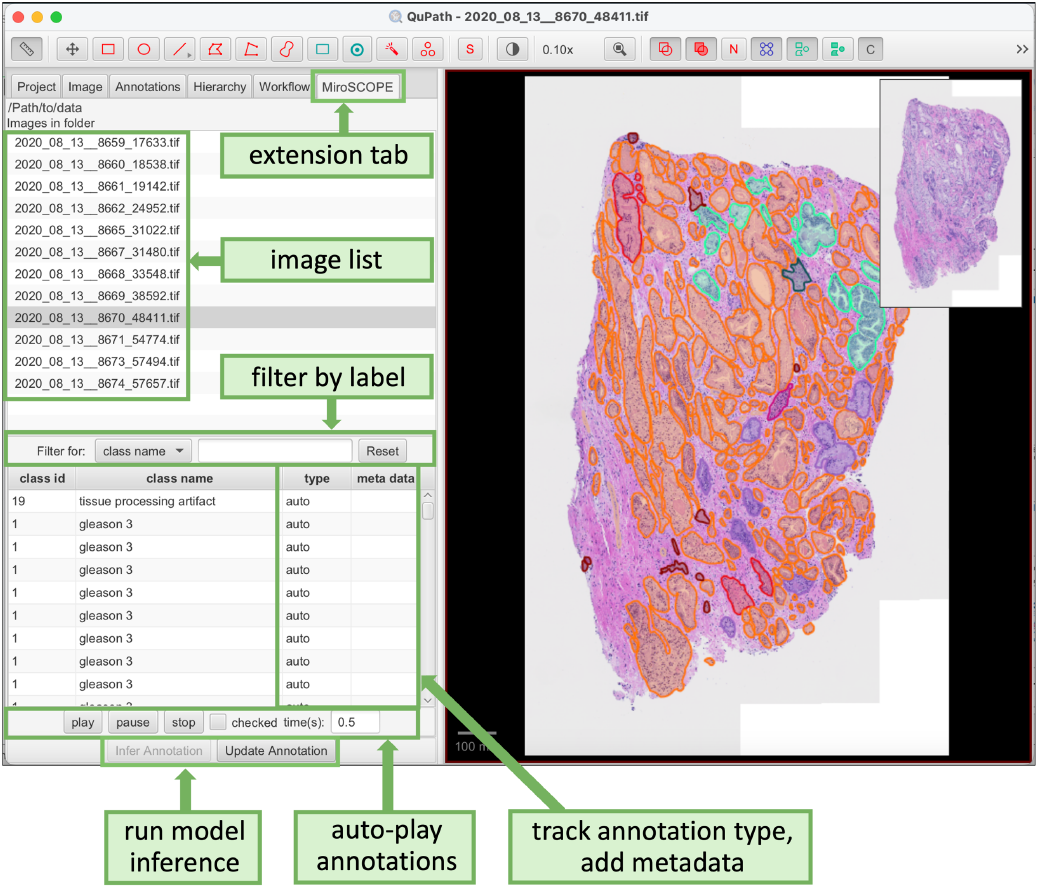
The QuPath-based MiroSCOPE platform offers enhanced user interfaces for annotating FTUs. Key features within the MiroSCOPE extension tab are highlighted.

The MiroSCOPE extension, organized under a tab UI element, allows users to initiate and control FTU segmentation and inference, identify and correct errors, and record the final annotations. Users can use this tab to select an image, and any existing annotations are then loaded into the annotation table. The annotation table serves as the main workspace, and actions are synchronized with the right-side image viewer. Each annotation is assigned a class ID, class name, type, and allows for the input of metadata, all of which can be edited in the table. Annotation type captures the degree of automation in generating the annotation, categorized using four labels:

- **auto**: An annotation created by an AI model.
- **auto_checked**: An annotation identified by an AI model and confirmed by a pathologist.
- **auto_edited**: An annotation identified by an AI model and edited by a pathologist.
- **manual**: An annotation created manually by a pathologist.

To streamline the review of FTU annotations, we also included an auto-play feature that automatically cycles through each FTU annotation. If an annotation is identified as requiring further inspection, it can be marked for later review. Annotations are saved in the GeoJSON format, with each FTU annotation identified by a unique UUID and associated metadata. This format is also used by the Python inference tool to facilitate the persistence and exchange of FTU annotations. All user actions are tracked to enable evaluation of changes in curation efficiency and provide feedback for model enhancements.

The QuPath extension-based platform provides a default trained AI model for prostate cancer FTU segmentation and classification. To extend the platform for AI models targeting other cancers or to support models fine-tuned for specific datasets, we have added UI features that allow users to configure the trained model and the associated FTU ID and class file.

### Fine-tuned segmentation foundation models accurately segment prostate tissue structures

When annotating FTUs, accurately defining the boundaries of each structure demands significant manual effort, making segmentation the most time-consuming step in the curation process. Encouraged by the recent performance gains by using a vision transformer (ViT) architecture in other domains, we applied the ViT-based Segment Anything Model (SAM)^24^ zero-shot to segment FTU structures in prostate cancer H&E. An evaluation of segmentation performance found an average Dice coefficient of 0.27. While SAM sometimes provided a useful starting point for segmentation correction, detecting boundaries of normal or early-stage well-defined structures reasonably well, it struggled with more disordered, higher-grade tumors.

After curating an initial set of annotated H&E images, we adopted a version of SAM that enables prompting by label and is readily fine-tunable. Given the limited number of samples and imbalanced class distributions, the model was first fine-tuned to perform foreground-background segmentation, considering FTU structures as foreground. After fine-tuning, the model reached an average Dice coefficient of 0.86. Following the curation of more images using the latest model checkpoint for assistance, a second fine-tuning was performed, achieving an average Dice coefficient of 0.91. A clear improvement in segmentation performance is observed at each fine-tuning (Figure 4). Since segmentation is the most time-consuming component of annotation, each fine-tuning offers substantial improvements for annotation throughput. Training times varied based on dataset size but were consistently completed within a few hours, a relatively small time investment for the resulting gains in efficiency.

**Figure 4.**
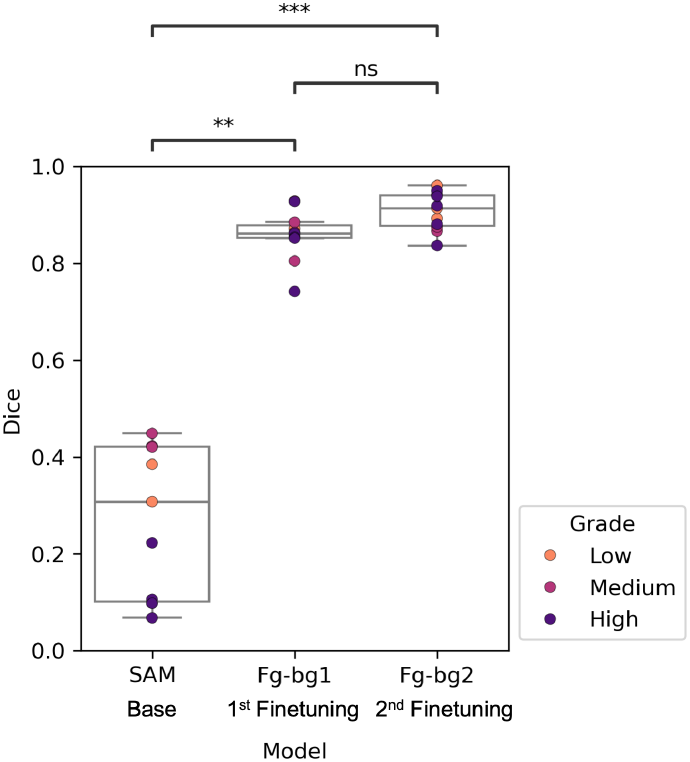
Evaluation of foreground-background model segmentation performance. Dice coefficient scores were calculated on a holdout dataset of prostate cancer H&Es (n=11) to evaluate the performance of SAM (vit-h) versus fine-tuned foreground-background SAM (vit-b) for segmenting FTUs. Checkpoints Fg-bg1 and Fg-bg2 of the foreground-background model were trained as more annotated H&E data was curated. Dice results are visualized as a box plot, where each image from the holdout set is represented by an overlayed point and coded by cancer grade (low: Gleason score <= 6, medium: Gleason score = 7, high: Gleason score > 7). Mean Dice scores for the SAM, Fg-bg1, and Fg-bg2 models were 0.27, 0.86, and 0.91 respectively. A pairwise Dunn’s test with Bonferroni correction between Dice scores for each model found significant differences between SAM and Fg-bg1 (p=0.0039) and SAM and Fg-bg2 (p= 5×10^−6^), but no significant difference between Fg-bg1 and Fg-bg2 (p=0.352).

### Fine-tuned segmentation foundation models accurately classify prostate tissue structures

Following the curation of 120 prostate cancer H&E samples, we fine-tuned a multiclass SAM model using the 17 FTU annotation labels defined in our controlled vocabulary. Model inference produces a pixel-wise classification, resulting in a semantic segmentation mask for the entire analyzed region. Visual observation of the results for unseen data shows that, while the model is more consistent for lower-grade cases, it regardless provides an excellent starting point for annotation correction (Figure 5A). Evaluation of the segmentation performance on the holdout set revealed that the fine-tuned multiclass model achieved an average Dice similarity coefficient of 0.81. While this score is lower than that achieved by the foreground-background model (Dice = 0.91), which is expected given the increased complexity of distinguishing 17 FTU classes plus background, it significantly outperforms the zero-shot SAM baseline (Dice = 0.27). We observed a similar trend regarding cancer grade as seen with original SAM, where lower and medium-grade cases generally yielded higher Dice scores compared to higher-grade cases, likely because tissue architecture is more preserved (Figure 5B)

**Figure 5.**
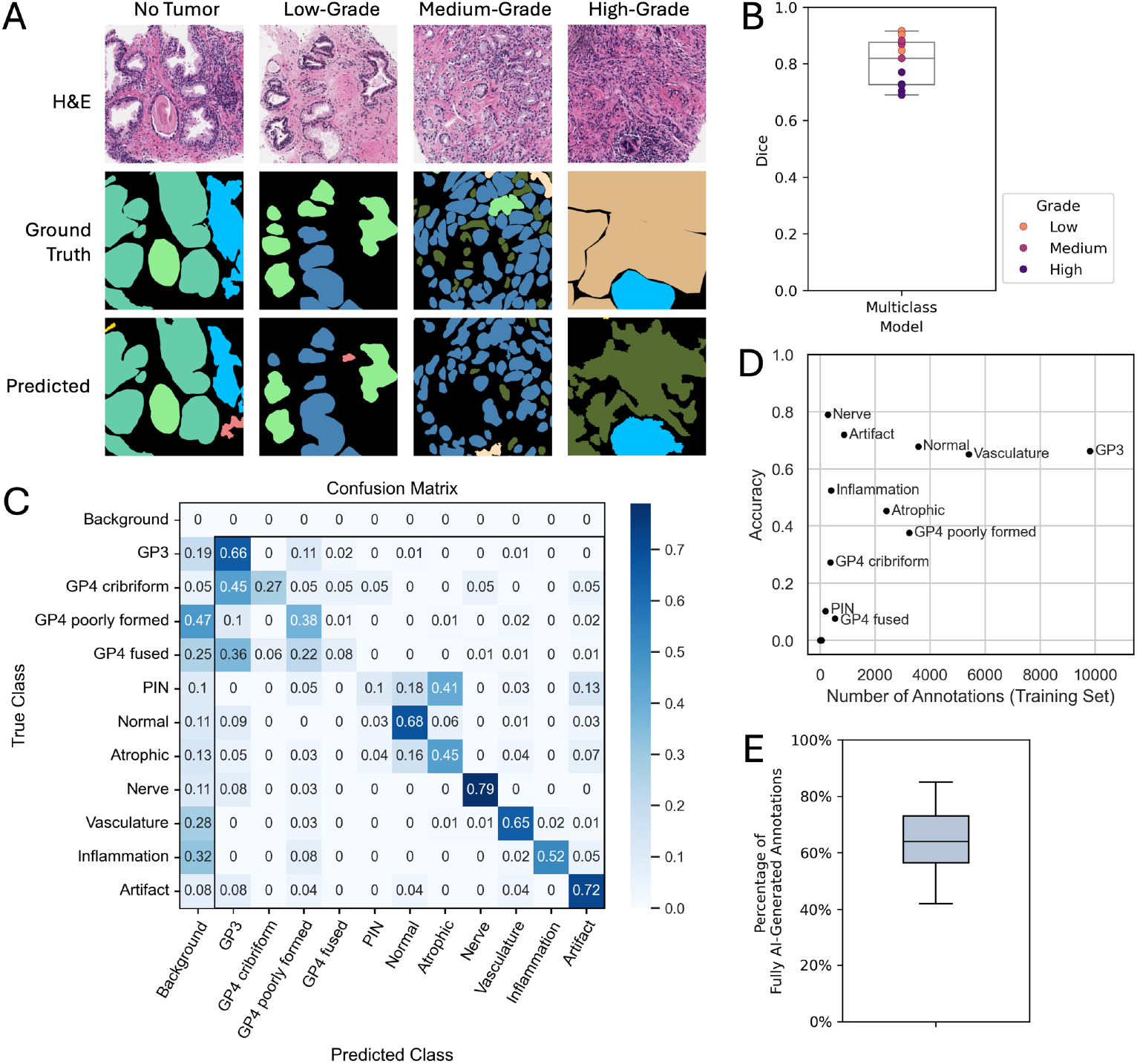
Evaluation of multiclass model segmentation and classification performance. The fine-tuned multiclass SAM (vit-b) model was trained to classify 17 categories of prostate cancer FTUs and evaluated on a holdout dataset of H&Es with prostatic acinar adenocarcinoma (n=11). To evaluate the practical improvements of using MiroSCOPE with the multiclass model, the annotation process of a separate efficiency evaluation dataset of 18 H&Es with prostatic acinar adenocarcinoma was tracked in detail. **A** Visualization of multiclass model predicted masks on holdout H&Es compared to ground truth masks for different cancer grades (no tumor, low-grade: Gleason score <= 6, medium-grade: Gleason score = 7, high-grade: Gleason score > 7). **B** Boxplot of Dice similarity coefficient scores calculated to evaluate segmentation performance. Each image from the holdout set is represented by an overlayed point and coded by cancer grade. The mean Dice across samples is 0.81. **C** Row-normalized confusion matrix visualizing classification accuracy for FTUs. Gleason pattern 5 labels and Gleason pattern 4 glomeruloids, which were underrepresented or absent from the training set, are not visualized (Supplementary Figure 1B). A full unnormalized confusion matrix can be found in Supplementary Figure 1A. GP: Gleason pattern; PIN: prostatic intraepithelial neoplasia. **D** Classification accuracy plotted against the number of annotations used for training, with each class represented by a labeled point. **E** Box plot of the percentage of annotations in each image from the efficiency evaluation dataset whose segmentations were fully generated by AI, requiring no manual intervention. The overall mean was 64%.

Assessing the classification performance of the fine-tuned multiclass model yielded an overall accuracy of 45.8%. Despite the challenge of classifying 17 distinct FTU categories, the model demonstrated strong discriminatory power for several key classes, far exceeding the 5.9% accuracy expected from random chance (Figure 5C). The model performed well at discriminating normal glands (68%), Gleason pattern 3 glands (66%), artifacts (72%), and stromal components such as nerves (79%) and vascular structures (65%). Performance was moderate for atrophic glands (45%), inflammation (52%), Gleason pattern 4 cribriform (27%), and Gleason pattern 4 poorly formed (38%). Classes corresponding to higher-grade Gleason patterns (other GP4 subtypes, GP5) and prostatic intraepithelial neoplasia (PIN) were largely underrepresented or absent in the training set, and consequently, their classification accuracies were close to a random classifier. Importantly, categories with many annotated samples generally achieved superior performance, and there were no classes which had many instances annotated but low classification accuracy (Figure 5D).

Since segmentation and classification performance are not entirely independent in semantic segmentation tasks, the evaluation of one must be understood in the context of the other. The most common source of misclassification across all classes was a failure to segment the structure, resulting in it being classified as background with a rate between 5% to 47% across all classes (Figure 5C). In effect, accuracy as a measure of discrimination between the 17 FTU classes is underestimated by the values presented. Another common source of error was misclassification as Gleason 3, most notably seen in Gleason 4 cribriform (45%) and Gleason 4 fused (36%), with lower rates for normal glands (9%), atrophic glands (5%), nerves (8%), and artifacts (8%). Since Gleason 3 has the highest representation in the training set of all classes, this likely indicates some bias of the model (Supplementary Figure 1B). However, another factor to consider is that diseases typically progress on a continuous trajectory, but grading systems define discrete boundaries along this progression to more easily characterize the disease. There are likely FTUs that exhibit mixed features of normal and Gleason 3, or Gleason 3 and Gleason 4, and misclassification between these groups may not necessarily reflect a failure of the model.

### MiroSCOPE greatly improves annotation efficiency

To assess gains in annotation efficiency when using MiroSCOPE with AI assistance, the annotation process for a set of 18 UCL AS Cohort images was tracked in detail. Evaluation focused on improvements in segmentation effort and time, as segmentation is the most time-consuming step in the annotation process. For the evaluation set, using AI assistance with the latest multiclass checkpoint was on average 3.18 more efficient than manual annotation alone. On average, 64% of annotations were fully AI generated without needing any manual segmentation creation or editing, saving an estimated 76 minutes of annotation time per image (Figure 5E; Supplementary Figure 4A). These efficiency measures varied greatly by image, with percentage of fully AI-generated annotations ranging from 42% to 85%, and estimated time saved ranging from 30 to 148 minutes. This is expected, as factors such as tissue size, complexity, and composition of cancer grades, together with model accuracy for different classes, can lead to variability in the annotation process (Supplementary Figure 4B).

### Miro-120 dataset: a high-quality FTU-level annotated dataset for the community

Using MiroSCOPE we curated Miro-120, a dataset consisting of 120 prostatic acinar adenocarcinoma H&E samples with detailed FTU annotations totaling 30,568 (Figure 7). We are offering Miro-120 as an openly available resource for the community, providing access to uniquely detailed H&E structure labeling to facilitate novel machine learning aims. To the best of our knowledge, this is the first prostate cancer H&E dataset with comprehensive structural annotations that include Gleason grade subpatterns and stromal components to be released. Samples in Miro-120 were sourced from two cohorts: 18 are radical prostatectomy specimens from the OHSU Biolibrary, and 102 are needle biopsy specimens from the CEDAR Biorepository. All patients were treatment-naïve with localized prostate cancer at the time of tissue collection. Annotations were curated using MiroSCOPE with AI assistance, and all annotations went through manual review and correction by an expert, making them highly reliable.

**Figure 7.**
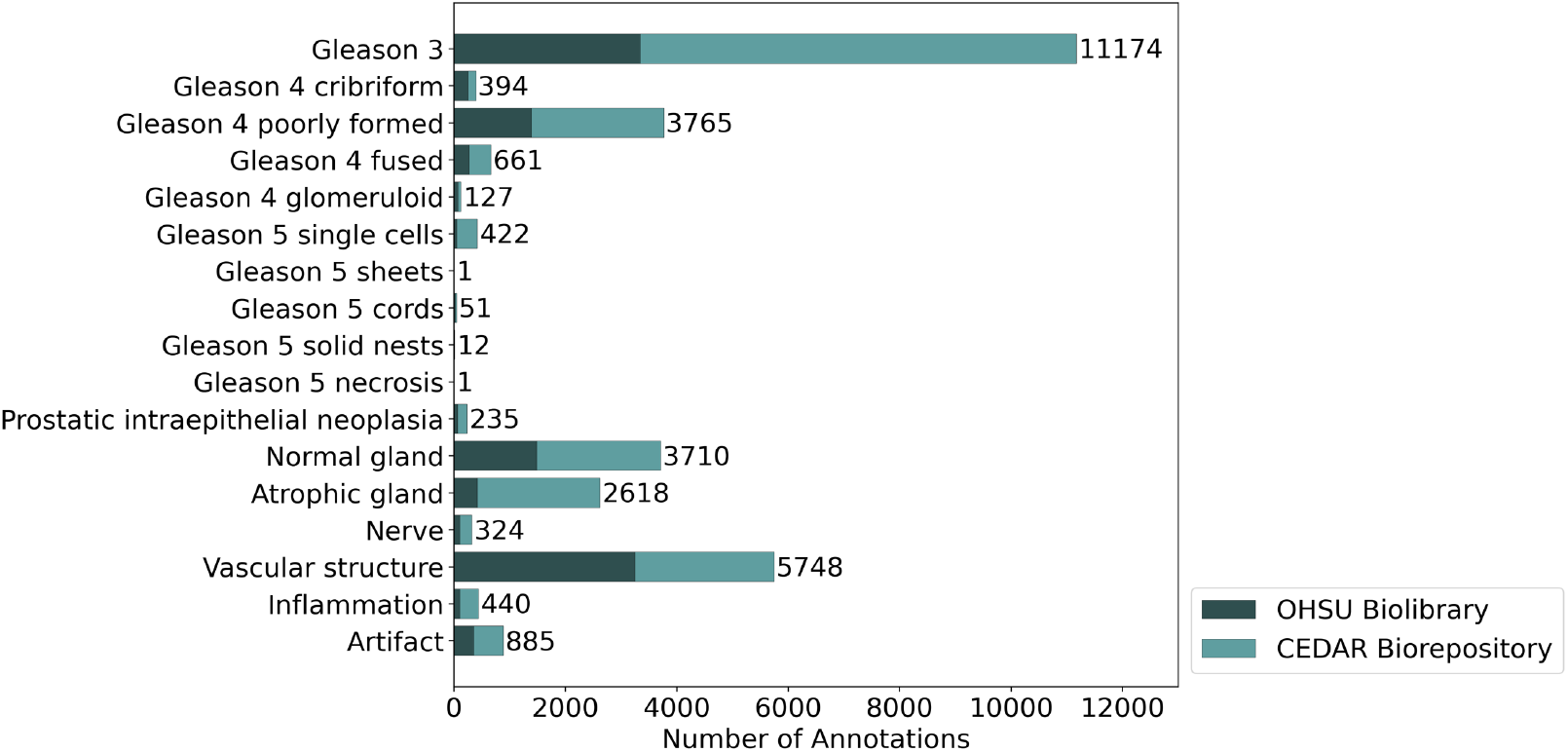
Annotation class counts for the Miro-120 dataset. The number of FTU annotations for each class label in the controlled vocabulary is visualized. The dataset consists of 120 prostatic acinar adenocarcinoma H&E samples sourced from two cohorts (OHSU Biolibrary (n=18); CEDAR Biorepository (n=102)) which are distinguished by color. The total annotation count across all classes is 30,568.

### The curation system can be deployed for other cancer types

While the curation study was piloted using prostatic acinar adenocarcinoma, the platform itself is generic. To evaluate transfer of the current approach to different cancer types, we used the fine-tuning capabilities offered within MiroSCOPE to adapt the model for annotating breast adenocarcinoma samples. The morphological similarities shared between adenocarcinomas make breast cancer a natural test for transfer learning. Following the structure of the controlled vocabulary defined for prostate FTUs, labels under non-tumoral, precursor, and tumoral categories were modified to reflect terms used for breast FTUs (Figure 8A). As this study on translating the curation system was limited in scope, terms are not exhaustive, and within non-glandular tumoral formations the label “solid” is used to collapse many high-grade patterns into a single term. Additionally, cribriform encompasses complex glandular structures, and benign tumors such as fibroadenoma were absent from the samples and not considered in the vocabulary.

**Figure 8.**
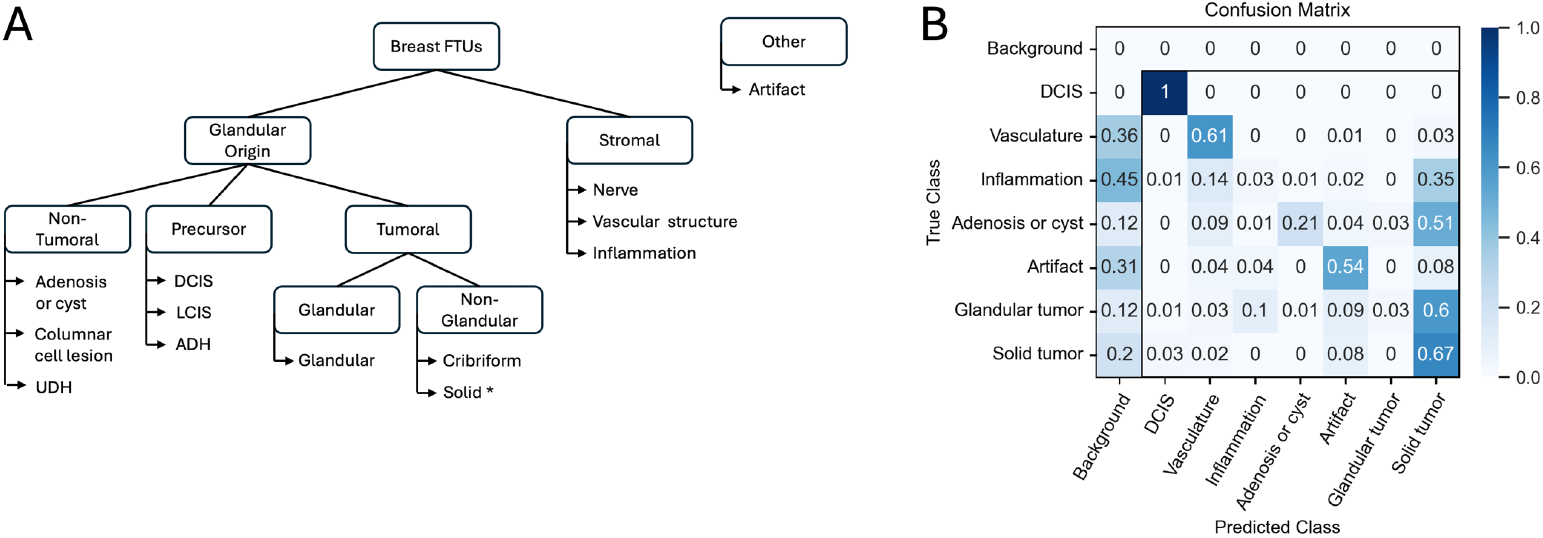
FTU categories and multiclass model classification performance for breast cancer. The FTU controlled vocabulary structure was adapted for breast cancer, then to assess translatability of the annotation system the AI annotation model was fine-tuned with a single TCGA prostate cancer H&E sample and classification performance was evaluated on another. **A** Hierarchical organization of breast FTU categories defined by the controlled vocabulary. *Solid tumor category encompasses multiple high-grade patterns (e.g. nests, cords, sheets). UDH: usual ductal hyperplasia; DCIS: ductal carcinoma in-situ; LCIS: lobular carcinoma in-situ, ADH: atypical ductal hyperplasia. **B** Row-normalized confusion matrix visualizing classification accuracy on one breast adenocarcinoma H&E sample from the TCGA-BRCA cohort following fine-tuning with another. Classes with less than 50 annotated instances for training are not shown (Supplementary Figure 2B); a full unnormalized confusion matrix can be found in Supplementary Figure 2A. DCIS: ductal carcinoma in-situ.

Two H&E samples from the TCGA-BRCA cohort with similar subtypes of breast adenocarcinoma were selected, one for fine-tuning a breast-specific annotation model and the other for evaluating model performance. To curate ground truth annotations for the first sample, the latest foreground-background model trained on prostate tissue was applied to accelerate annotation correction. As a baseline, the Fg-bg2 model checkpoint which had only seen prostate images achieved a Dice of 0.68 on the test image. To compare approaches, both a foreground-background model and a multiclass model were trained using the breast H&E sample. The foreground-background model was trained from the Fg-bg2 model checkpoint and reached a Dice of 0.77. The multiclass model was trained from the latest multiclass checkpoint and reached a Dice of 0.68 and overall accuracy of 49.7%.

We observed successful classification of ductal carcinoma in-situ (100%), solid tumors (67%), vascular structures (61%), and artifacts (54%) (Figure 8B). Similarly to the multiclass model trained on prostate images, failure to segment the structure and instead labeling it as background was a large source of misclassification, particularly for vascular structures (36%) and inflammation (45%). Additionally, a clear bias is observed for classifying solid tumors, with many false positives in other classes indicating low specificity. This is likely because solid tumors were over-represented in the training image, highlighting the risk of fine-tuning with just one sample - a situation that is hard to avoid at the start of curation (Supplementary Figure 2B).

## Discussion

In this work, we present MiroSCOPE, an AI-driven, human-in-the-loop annotation platform centered around functional tissue units, designed to address limitations in current digital pathology approaches. MiroSCOPE is a QuPath extension that integrates a fine-tunable AI model for annotation, facilitating large-scale, expert-driven annotation of FTUs and bridging the gap between cellular detail and tissue-level architecture. We demonstrated its utility by annotating over 71,900 FTUs in prostate cancer and showed its ready translation to breast cancer.

Tissue FTUs represent the fundamental building blocks of an organ, where intercellular coordination enables critical functions. They offer a biologically meaningful scale for tissue analysis, connecting cellular detail and overall tissue architecture, and offering a window into tumor progression and invasion. In diseases like cancer, the progressive breakdown of this coordination is often most clearly visualized as morphological disorganization at the FTU level^25^. Analyzing tissues through the lens of FTUs offers a more biologically grounded perspective, analogous to how modern natural language processing moved beyond word frequencies (“bag-of-words”) to understand contextual relationships across sentences and documents. Integrative analysis of FTU features holds promise for enhancing understanding of tumor progression and informing multi-cellular disease modeling.

Analysis at the FTU level is complimentary to tissue analysis approaches that operate at different granularities, such as cell-level or tile-level: While a tile-based model might predict a high-risk score for a region, FTU annotations can reveal if this is due to the presence of a certain type of tumor glands, specific stromal reactions, nerve invasion, or other structurally defined features. This allows for validation and interpretation of tile-based predictions in a language pathologists can interpret. However, unlike cell- or tile-based methods which benefit from existing datasets or established self-supervised techniques respectively, FTU-level analysis has been hindered by the lack of large-scale, detailed annotations. For this reason, methods incorporating FTU information during tissue analysis are sparse^15,26^. MiroSCOPE is designed to overcome this specific challenge by enabling efficient, large-scale curation of FTU annotations.

Standardizing the language for annotating FTUs provides a connection between modern AI approaches and clinical literature. A critical component of any annotation system is a standardized ontology with clarity, comprehensiveness and comprehensibility^27^. While ontologies for healthy FTUs and cell types are actively developing^28–30^, standardized descriptions for FTUs in disease states are largely lacking, despite extensive documentation in pathology literature. For this study, we developed a practical, 17-term controlled vocabulary for prostatic acinar adenocarcinoma FTUs, grounded in the Gleason system and encompassing key non-tumoral and stromal elements. This vocabulary proved readily adaptable to breast adenocarcinoma by substituting relevant terms.

The controlled vocabulary presented in the study is intended as an initial step. Future efforts should focus on developing comprehensive, shared FTU ontologies that formally connect to existing resources like the Cell Ontology (CL)^29^ and UBERON^30^. Such an ontology should extend beyond hierarchical classification to capture cross-cutting morphological features (e.g., “foamy,” “microcystic,” corresponding to unusual but recognized subtypes we excluded initially for clarity) and pathological processes (e.g., “inflammation,” “loss of basal membrane”). This would bridge classical pathology knowledge with modern AI, enhance interpretability, and facilitate comparative studies across organs and diseases. Standardizing terminology is essential for building the large, reusable datasets needed to train robust FTU-level models.

To enable large-scale FTU annotation, MiroSCOPE combines a fine-tunable AI annotation model, UI features for curation, and a controlled vocabulary, creating a HITL system within the widely adopted QuPath environment. While general-purpose segmentation models like SAM provide a zero-shot starting point, we found their performance on complex pathology images, especially higher-grade tumors, can be limited. Additionally, existing FTU annotation models cannot reliably generalize to new datasets. Our HITL workflow allows pathologists to efficiently correct initial AI segmentation and classification, generating data to iteratively fine-tune the model. This cycle substantially accelerates the creation of high-quality annotations, and a specialized, high-performing model tailored to the specific task and dataset. In the future, we plan to expand on these features to offer collaboration options, better versioning and tracking, expanded controlled vocabularies and expanding AI models into other tumor types.

By using a transformer-based architecture for FTU segmentation and classification, we achieved sufficient accuracy to initiate a HITL workflow. A limited number of automated FTU segmentation algorithms exist, and most fail to cover all FTU presentations, including tumoral and stromal^13–16^. MiroSCOPE’s annotation workflow was employed to capture the entire spectrum of the prostate cancer disease state, in addition to features present in the surrounding stroma. As discussed in the results section, assessing the true performance of the AI model is challenging due to various modes of failure, class representation biases and high interobserver variability and inherent ambiguity in “ground truth” classifications. Importantly, we observed that the initial segmentation and classification performance achieved using MiroSCOPE was sufficient to enable a HITL workflow to iteratively improve the annotation rate and model performance. While it is challenging to measure generalized efficiency gains due to variations in tissue factors like size and cancer grade composition, our assessment on a subset of images found the final multiclass prostate model made the annotation process on average 3.18 more efficient, significantly speeding up the process. Additionally, models trained using prostate cancer images showed translation to breast cancer after fine-tuning with one image, demonstrating the applicability to other cancer types.

Using MiroSCOPE, we curated annotated prostate cancer images with segmentations and labeling of all tissue structures. We publicly release Miro-120, a high-quality dataset of 120 prostatic acinar adenocarcinoma H&E with 30,568 annotations that comprehensively cover all non-tumoral, tumoral (with subpatterns), and stromal features of the tissue, offering a unique level of detail which is not present in existing datasets^31^. This dataset can be used by the community as a resource for FTU-level machine learning studies that incorporate Gleason subpatterns and stromal features of tissue structure.

In the future, MiroSCOPE can act as a common platform for building FTU datasets and analyzing them with modern deep-learning methodologies. Common data marketplaces such as the Protein Data Bank (PDB)^32^, the database of Genotypes and Phenotypes (dbGAP)^33^, or The Cancer Genome Atlas (TCGA)^34^ project allow computational biologists to develop algorithms, servers, resources and tools once, but then apply them to a large number of diseases and datasets. Decades after their inception, these common resources remain highly central for their communities and are used every day for testing, evaluating, and applying algorithms. A similar opportunity exists here, where a common resource for well annotated pathology images can spur algorithm development to better understand the disease microenvironment across a large context. Topics we can address include disease prognosis, cell-cell dynamics within FTUs, 3D reconstruction across tissue sections, and higher-level interactions between FTUs and immune and neural processes. MiroSCOPE also holds the potential to facilitate clinically driven aims related to interobserver variability feedback and detection of border cases with high classification error.

We have demonstrated that it is possible to accelerate FTU curation with an iterative human-in-the-loop system even with modest datasets and limited expert annotation. There are substantial shared features between different FTUs, such as basal membrane, vasculature, or stroma. As we expand the curated datasets to other tumors and diseases, we expect the AI models to increasingly improve at understanding these common components. Our results indicate that modern ViT architectures can effectively transfer these learned features between disease contexts, further accelerating model training.

## Methods

### Data

Whole slide image H&E samples of prostatic acinar adenocarcinoma were sourced from three unique cohorts which are referred to as OHSU Biolibrary, CEDAR Biorepository, and UCL AS Cohort, making 184 images total. The OHSU Biolibrary data refers to 18 radical prostatectomy samples from the Oregon Health and Science University (OHSU) Knight Cancer Institute (KCI) Biolibrary (IRB#4918). The CEDAR Biorepository data refers to 102 needle biopsy samples obtained from the Cancer Early Detection Advanced Research (CEDAR) Specimen and Data Repository (IRB#18048). Samples of the OHSU Biolibrary and CEDAR Biorepository are collected from clinics across OHSU following informed consent and in accordance with OHSU institutional review boards. The UCL AS Cohort refers to 64 needle biopsy images which were collected at University College London (UCL) hospitals for Active Surveillance (AS). For the current study, all samples were de-identified, and work was not considered human subjects research.

### Software development

We followed the software architecture of the MONAI Label plugin for QuPath (https://github.com/Project-MONAI/MONAILabel/tree/main/plugins/qupath) to develop our QuPath-based MiroSCOPE platform for AI-assisted annotation curation. To create the Java-based QuPath user interfaces, we forked the main branch of the qupath codebase (as of June 11, 2024), updated the default Java version to 21, and adopted the Gradle-based build scripts in the original QuPath codebase. We developed the Java-based code using IntelliJ IDEA (Community Edition, version 2024.1.4). For offering an AI annotation model for automated FTU segmentation and classification, we utilized MONAI Label’s Python-based architecture. Development was conducted in Python 3.9 within a Conda environment, using VS Code as the primary IDE. The Java code is hosted at https://github.com/ohsu-cedar-comp-hub/qupath_monaillabel_plugin the Python code at https://github.com/ohsu-cedar-comp-hub/monailabel_cedar_app.

Initial iterations of the HITL system used the image viewer napari^35^ as the interface for annotation correction. A custom napari plugin with UI for managing annotations was developed, and many features were transferred to the current MiroSCOPE QuPath extension.

### Training details

To demonstrate using MiroSCOPE for HITL curation, we fine-tuned the annotation model as ground truth annotation data was generated. Images were divided into tiles of dimension 1024×1024 as input for training, with 20% randomly assigned to the validation set. All training used AdamW optimization^36^, generic data augmentations, and identical parameters including a learning rate of 1×10^−4^, regularization weight of 1×10^−5^, and momentum of 0.99. Final model weights were chosen based on best validation accuracy. Either a Nvidia Tesla V100 32GB DRAM GPU or Nvidia A40 44GB DRAM GPU were used for computation. The curation system was primarily demonstrated with internal prostate cancer samples, but translatability to other cancer types was also piloted with breast cancer data from the TCGA-BRCA dataset. For fine-tuning with prostate cases, the Fg-bg1 checkpoint was trained using 10 OHSU Biolibrary samples, the Fg-bg2 checkpoint was trained using 16 OHSU Biolibrary samples, the multiclass model was trained using 18 OHSU Biolibrary and 91 CEDAR Biorepository samples. For fine-tuning with breast cases, a single TCGA-BRCA sample (TCGA-A2-A259) was used.

### Model evaluation

A holdout test set of 11 images from the Biorepository cohort which have representation across cancer grades was used for all prostate-specific model evaluations. For running inference for original SAM (vit-h), images were divided into 1200×1200 tiles with 5% overlap before running inference on each, then borders were resolved during restitching. For running inference with class promptable SAM (vit-b), a sliding window approach using patches of size 1024×1024 was used to create a seamless annotation mask. All masks underwent post processing to fill holes and filter objects below a pixel area threshold of 2000. To evaluate segmentation performance, the Dice similarity coefficient^37^ was computed between the ground truth mask and predicted mask (Figure 9A). For the multiclass model, the Dice was calculated in the same manner without considering class, and accuracy was determined by comparing the class assignment for the center pixel of the ground truth annotation with the same pixel of the predicted mask (Figure 9B). For breast-specific model evaluations, a single TCGA-BRCA sample (TCGA-A2-A0ES) was used, and the segmentation and multiclass model performance analyses were performed in the same way.

**Figure 9.**
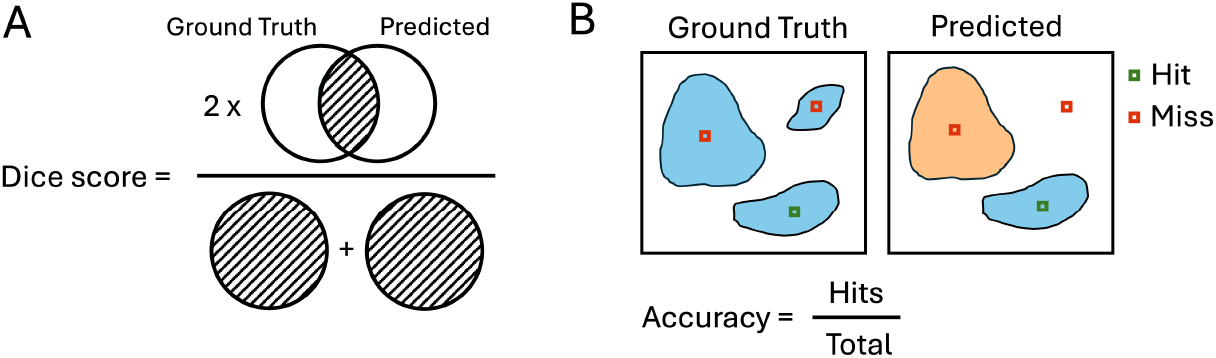
Evaluation metrics for segmentation and classification. **A** Segmentation performance is measured using the Dice similarity coefficient, which is calculated by taking twice the overlap between the ground truth and predicted masks and dividing it by the total area between both. **B** Classification performance is measured by determining whether the center pixel of a ground truth annotation is a “hit” or “miss” when compared to the same pixel of the predicted mask, then accuracy is calculated to be the number of hits over the total number of annotations.

### Efficiency evaluation

To assess gains in efficiency when using the platform, 18 images from the UCL AS Cohort were annotated in MiroSCOPE using AI assistance while all actions were tracked in detail. For each image, the number of manual annotations (*A*_*m*_), which described annotations that were manually segmented or AI-generated annotations that were manually edited, and the total number of annotations (*A*_*t*_), were recorded. The average manual segmentation time (*t*) was estimated at 10s per annotation (Supplementary Figure 3). Calculations of percentage of manual annotations, efficiency gain, percentage of annotations fully generated by AI, and time saved were made for each image, then averaged to estimate more generalized gains of using MiroSCOPE across varied images.

**percentage of manual annotations** = 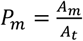

**efficiency gains** = 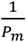

**percentage of annotations fully generated by AI** = 1 − *P*_*m*_

**time saved** = *t* × (*A*_*t*_ − *A*_*m*_)

### Statistics

To measure the significance of the increase in segmentation performance with each fine-tuning, the Kruskal-Wallis test was performed for the holdout test Dice coefficient scores across the three models (SAM, Fg-bg1, Fg-bg2) and found a p-value of 6.7×10^−6^. To investigate more specifically which models have significant interactions, a pairwise Dunn’s test was done for each grouping, and Bonferroni correction was applied to adjust p values.

## Data Availability

The Miro-120 dataset, and all other data and supporting files, are available at https://www.synapse.org/Synapse:syn66298514/wiki/.

## Code Availability

The MiroSCOPE extension is open-source and available for download at https://github.com/ohsu-cedar-comp-hub/qupath_monaillabel_plugin. The MONAI Label back-end to support AI annotation within the platform, as well as scripts for model inference and fine-tuning, are available at https://github.com/ohsu-cedar-comp-hub/monailabel_cedar_app.

## Acknowledgments

This project was supported by funding from the Cancer Early Detection Advanced Research Center (CEDAR Project ID# Full 2023-1748) at Oregon Health & Science University, Knight Cancer Institute. S. Ece Eksi was supported by funding from the International Alliance for Cancer Early Detection (ACED) (ACED 2022-1508) at Oregon Health & Science University, Knight Cancer Institute. Mark Emberton receives research support from the United Kingdom’s National Institute of Health Research (NIHR) UCLH/UCL Biomedical Research Centre. The research reported in this publication used computational infrastructure supported by the Office of Research Infrastructure Programs, Office of the Director, of the National Institutes of Health under Award Number S10OD034224. The content is solely the responsibility of the authors and does not necessarily represent the official views of the National Institutes of Health.

## Author Contributions

**M.R. Fenner:** Methodology, investigation, data curation, formal analysis, validation, visualization, project administration, writing – original draft, writing – review and editing. **S. Sevim:** Data curation, resources, writing – review and editing. **G. Wu:** Software, formal analysis, writing – review and editing. **D. Beavers:** Software, writing – review and editing. **P. Guo:** Methodology, investigation, validation. **Y. Tang:** Software. **C.Z. Eddy:** Software, investigation, writing – review and editing. **K. Ait-Ahmad:** Investigation. **T. Rice-Stitt:** Data curation. **G. Thomas:** Resources. **M.J. Kuykendall:** Resources. **V. Stavrinides**: Resources. **M. Emberton:** Resources. **D. Xu:** Supervision. **X. Song:** Conceptualization, supervision, funding acquisition. **S.E. Eksi:** Conceptualization, supervision, funding acquisition, writing – review and editing. **E. Demir:** Conceptualization, supervision, project administration, funding acquisition, writing – original draft, writing – review and editing.

## References

1. de Bono, B., Grenon, P., Baldock, R. & Hunter, P. Functional tissue units and their primary tissue motifs in multi-scale physiology. J Biomed Semantics 4, 22 (2013).

2. Xu, H. et al. A whole-slide foundation model for digital pathology from real-world data. Nature 630, 181–188 (2024).

3. Chen, R. J. et al. Towards a general-purpose foundation model for computational pathology. Nat Med 30, 850–862 (2024).

4. Jiang, X. et al. End-to-end prognostication in colorectal cancer by deep learning: a retrospective, multicentre study. The Lancet Digital Health 6, e33–e43 (2024).

5. Saillard, C. et al. Predicting Survival After Hepatocellular Carcinoma Resection Using Deep Learning on Histological Slides. Hepatology 72, 2000 (2020).

6. Hu, Y. et al. Unsupervised and supervised discovery of tissue cellular neighborhoods from cell phenotypes. Nat Methods 21, 267–278 (2024).

7. Schürch, C. M. et al. Coordinated Cellular Neighborhoods Orchestrate Antitumoral Immunity at the Colorectal Cancer Invasive Front. Cell 182, 1341-1359.e19 (2020).

8. Black, S. et al. CODEX multiplexed tissue imaging with DNA-conjugated antibodies. Nat Protoc 16, 3802–3835 (2021).

9. Peng, H. et al. Multiplex immunofluorescence and single-cell transcriptomic profiling reveal the spatial cell interaction networks in the non-small cell lung cancer microenvironment. Clin Transl Med 13, e1155 (2023).

10. Salvi, M. et al. A hybrid deep learning approach for gland segmentation in prostate histopathological images. Artif Intell Med 115, 102076 (2021).

11. Zhang, H. et al. Masked Image Modeling Meets Self-Distillation: A Transformer-Based Prostate Gland Segmentation Framework for Pathology Slides. Cancers (Basel) 16, 3897 (2024).

12. Qiu, Y. et al. Automatic Prostate Gleason Grading Using Pyramid Semantic Parsing Network in Digital Histopathology. Front Oncol 12, 772403 (2022).

13. Jain, Y. et al. Segmenting functional tissue units across human organs using community-driven development of generalizable machine learning algorithms. Nat Commun 14, 4656 (2023).

14. Godwin, L. L. et al. Robust and generalizable segmentation of human functional tissue units. 2021.11.09.467810 Preprint at 10.1101/2021.11.09.467810 (2021).

15. Ferrero, A. et al. HistoEM: A Pathologist-Guided and Explainable Workflow Using Histogram Embedding for Gland Classification. Modern Pathology 37, 100447 (2024).

16. Barmpoutis, P. et al. A digital pathology workflow for the segmentation and classification of gastric glands: Study of gastric atrophy and intestinal metaplasia cases. PLoS One 17, e0275232 (2022).

17. Ochi, M., Komura, D. & Ishikawa, S. Pathology Foundation Models. JMA J 8, 121–130 (2025).

18. Maximum androgen blockade in advanced prostate cancer: an overview of 22 randomised trials with 3283 deaths in 5710 patients. Prostate Cancer Trialists’ Collaborative Group. Lancet 346, 265–269 (1995).

19. Gleason, D. F. Classification of prostatic carcinomas. Cancer Chemother Rep 50, 125–128 (1966).

20. Epstein, J. I., Allsbrook, W. C., Amin, M. B. & Egevad, L. L. The 2005 International Society of Urological Pathology (ISUP) Consensus Conference on Gleason Grading of Prostatic Carcinoma. 29, (2005).

21. Epstein, J. I. et al. The 2014 International Society of Urological Pathology (ISUP) Consensus Conference on Gleason Grading of Prostatic Carcinoma: Definition of Grading Patterns and Proposal for a New Grading System. Am J Surg Pathol 40, 244–252 (2016).

22. Bankhead, P. et al. QuPath: Open source software for digital pathology image analysis. Sci Rep 7, 16878 (2017).

23. Diaz-Pinto, A. et al. MONAI Label: A framework for AI-assisted interactive labeling of 3D medical images. Medical Image Analysis 95, 103207 (2024).

24. Kirillov, A. et al. Segment Anything. Preprint at https://arxiv.org/abs/2304.02643v1 (2023).

25. Hanahan, D. & Weinberg, R. A. Hallmarks of cancer: the next generation. Cell 144, 646–674 (2011).

26. Sobral-Leite, M. et al. A morphometric signature to identify ductal carcinoma in situ with a low risk of progression. npj Precis. Onc. 9, 1–13 (2025).

27. Demir, E. et al. An ontology for collaborative construction and analysis of cellular pathways. Bioinformatics 20, 349–356 (2004).

28. Osumi-Sutherland, D. et al. Cell type ontologies of the Human Cell Atlas. Nat Cell Biol 23, 1129– 1135 (2021).

29. Diehl, A. D. et al. The Cell Ontology 2016: enhanced content, modularization, and ontology interoperability. J Biomed Semantics 7, 44 (2016).

30. Mungall, C. J., Torniai, C., Gkoutos, G. V., Lewis, S. E. & Haendel, M. A. Uberon, an integrative multi-species anatomy ontology. Genome Biology 13, R5 (2012).

31. Zhu, L. et al. Harnessing artificial intelligence for prostate cancer management. Cell Reports Medicine 5, 101506 (2024).

32. wwPDB consortium. Protein Data Bank: the single global archive for 3D macromolecular structure data. Nucleic Acids Research 47, D520–D528 (2019).

33. Mailman, M. D. et al. The NCBI dbGaP database of genotypes and phenotypes. Nat Genet 39, 1181–1186 (2007).

34. Weinstein, J. N. et al. The Cancer Genome Atlas Pan-Cancer analysis project. Nat Genet 45, 1113– 1120 (2013).

35. Chiu, C.-L., Clack, N., & the napari community. napari: a Python Multi-Dimensional Image Viewer Platform for the Research Community. Microscopy and Microanalysis 28, 1576–1577 (2022).

36. Loshchilov, I. & Hutter, F. Decoupled Weight Decay Regularization. Preprint at 10.48550/arXiv.1711.05101 (2019).

37. Dice, L. R. Measures of the Amount of Ecologic Association Between Species. Ecology 26, 297–302 (1945).

