## Supplementary Figures for "MiroSCOPE: An AI-driven digital pathology platform for annotating functional tissue units"

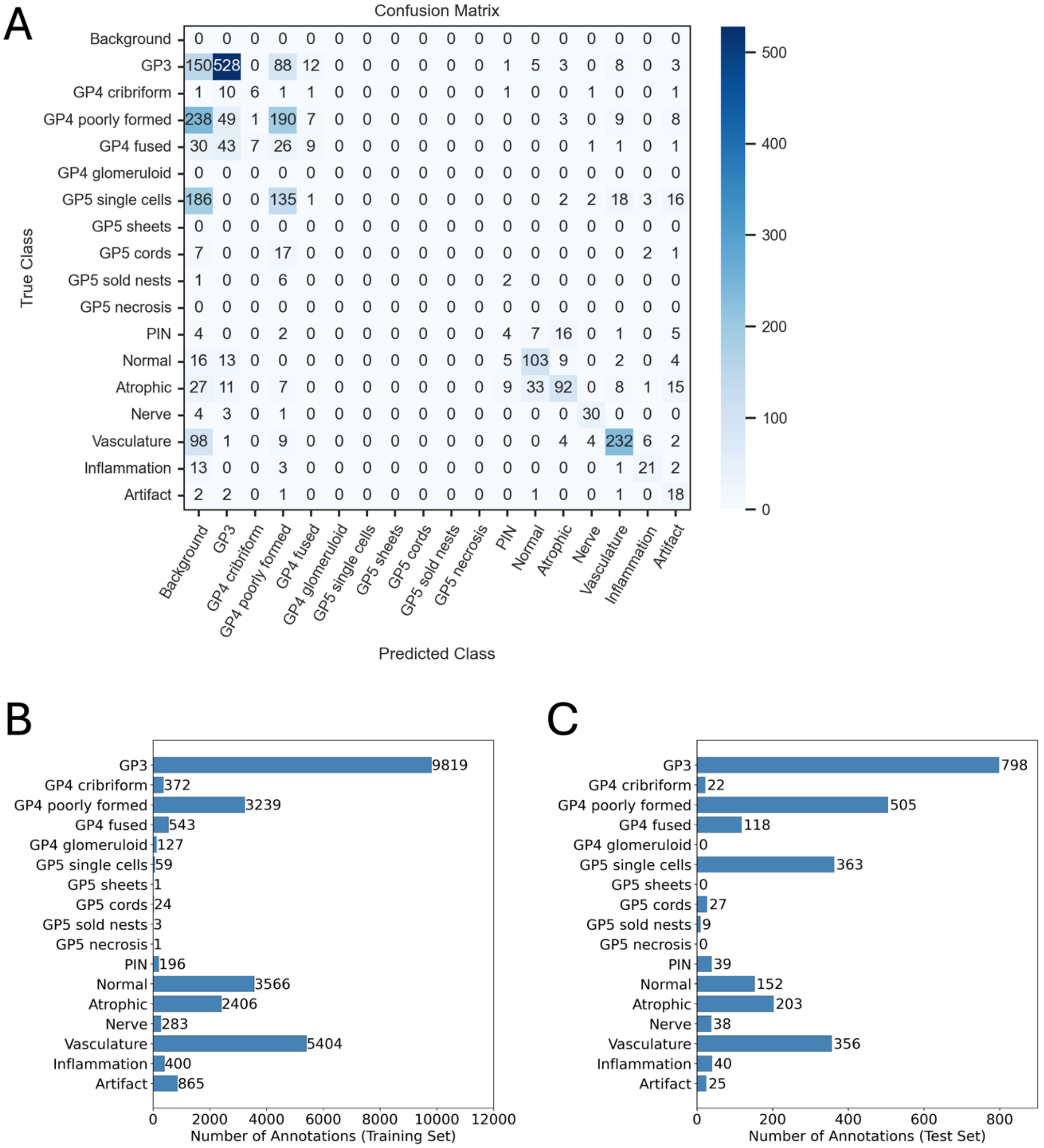


**Supplementary Figure 1. Multiclass prostate model data and evaluation details.** The fine-tuned multiclass SAM (vit-b) model was trained to classify 17 categories of prostatic acinar adenocarcinoma FTUs with a training set of 109 H&Es and evaluated on a holdout dataset of 11 H&Es. **A** Confusion matrix showing raw counts for classification on the holdout test set. **B** Training set (includes training and validation data) class distributions. **C** Holdout test set class distributions. GP: Gleason pattern; PIN: prostatic intraepithelial neoplasia.

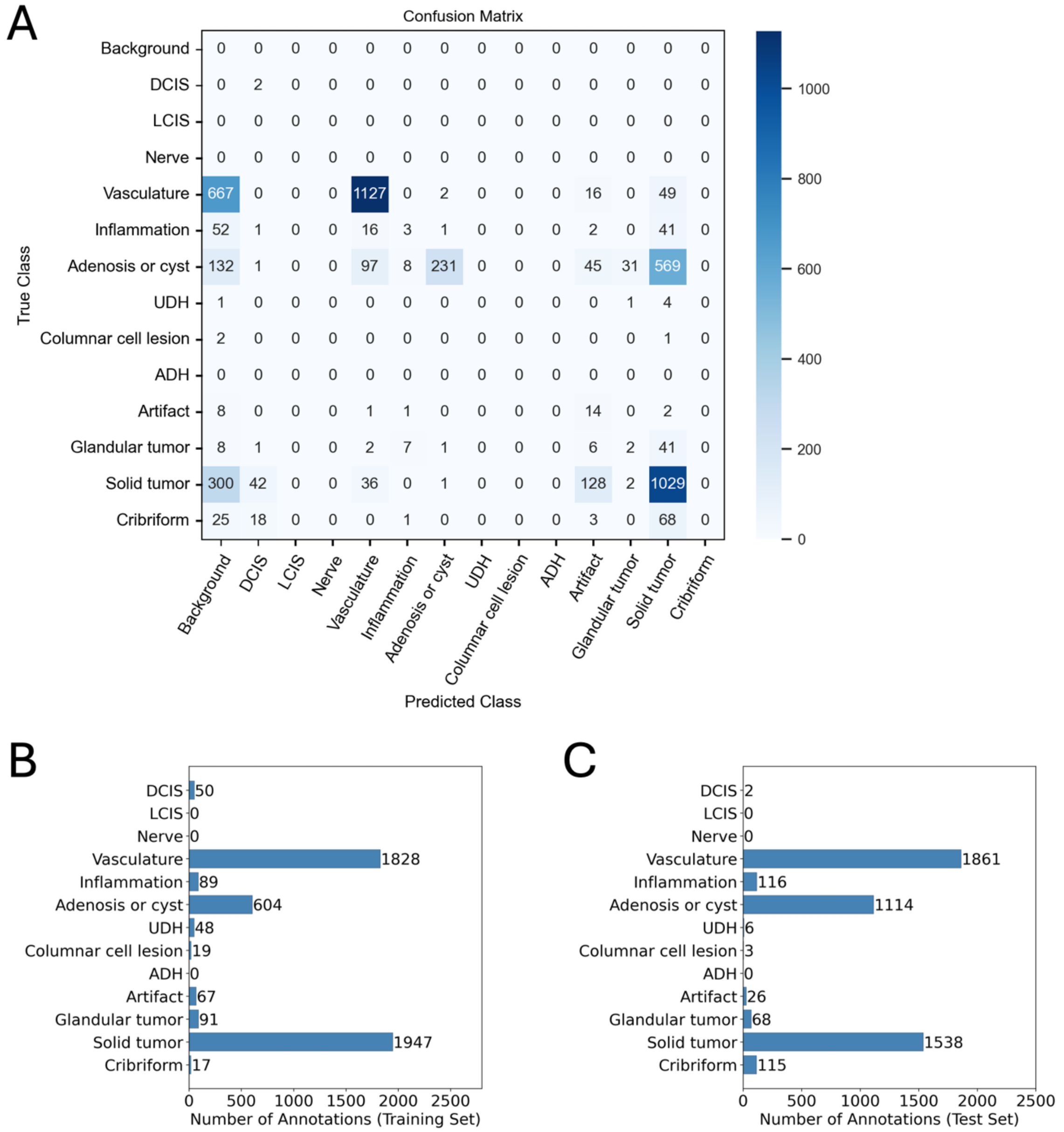


**Supplementary Figure 2. Multiclass breast model data and evaluation details.** The multiclass SAM (vit-b) model was fine-tuned with a single TCGA prostate cancer H&E sample and classification performance was evaluated on a separate TCGA prostate cancer H&E sample. **A** Confusion matrix showing raw counts for classification on the holdout test set. **B** Training set (includes training and validation data) class distributions. **C** Holdout test set class distributions. DCIS: ductal carcinoma in-situ; LCIS: lobular carcinoma in-situ; UDH: usual ductal hyperplasia; ADH: and atypical ductal hyperplasia.

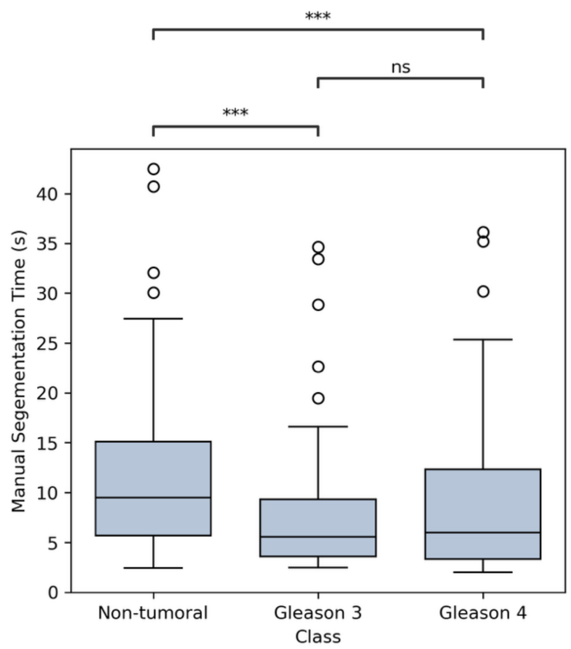


**Supplementary Figure 3. Manual annotation time comparison between classes.** A pathologist manually annotated instances of non-tumoral, Gleason 3, and Gleason 4 FTUs in MiroSCOPE without AI assistance while actions were tracked, and 100 instances of each were sampled for a comparison. Outliers over 45 s, which were determined to measure downtime rather than annotation time, were excluded. The box plot shows manual segmentation time (seconds) for sampled annotations, compared between cancer grade groups. Mean segmentation time is 12.60 s, 7.44 s, and 10.18 s for non-tumoral, Gleason 3, and Gleason 4 respectively, with an overall mean of 10.07 s. The difference in segmentation time between grade groups is significant by a Kruskal-Wallis test ($p=6.67 \times 10^{-6}$). A pairwise Dunn's test with Bonferroni correction between segmentation time for each grade group found significant differences between non-tumoral and Gleason 3 ($p=2.2 \times 10^{-5}$) and Non-tumoral and Gleason 4 ($p= 2.67 \times 10^{-4}$), but no significant difference between Gleason 3 and Gleason 4 (p=1).

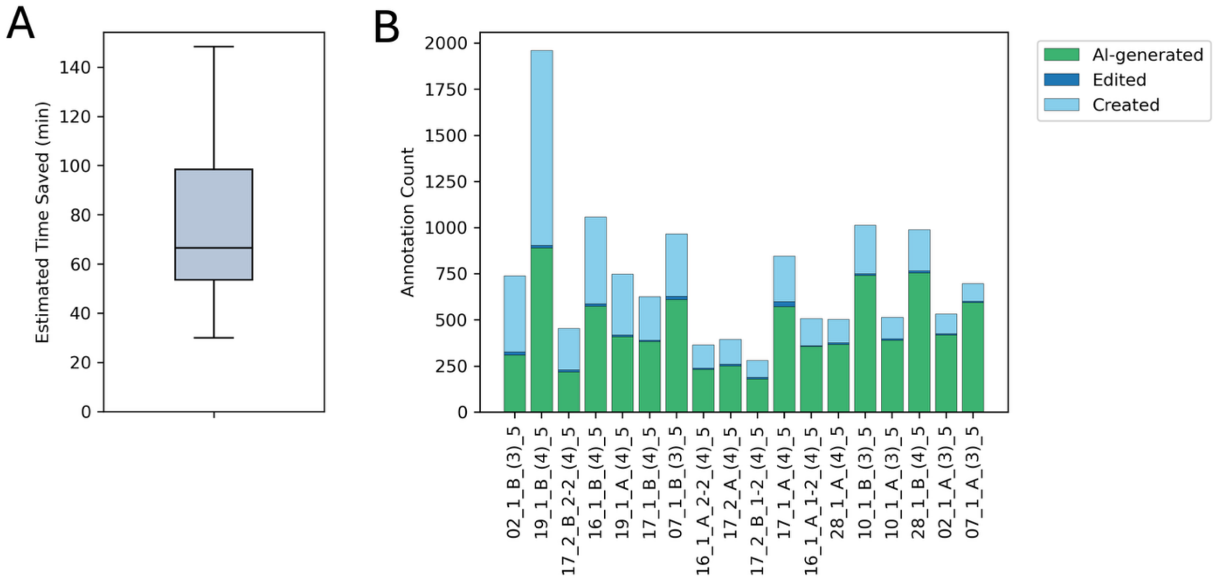


**Supplementary Figure 4. Efficiency gains when using MiroSCOPE for annotation.** To estimate measures of efficiency, the annotation process of 18 images from the UCL AS Cohort was tracked in detail. **A** Box plot of the estimated time saved by using AI assistance during annotation for each image, calculated assuming an average manual segmentation time of 10s per annotation (Supplementary Figure 3). The mean time saved per image was 76 minutes. **B** Counts of AI-generated, edited, or created annotations for each of the UCL AS Cohort samples used for efficiency evaluations, ordered left to right by percentage of AI-generated annotations from lowest to largest. Edited annotations are AI-generated annotations that are manually edited, and created annotations are manually segmented.
